# DIS3L, cytoplasmic exosome catalytic subunit, is essential for development but not cell viability in mice

**DOI:** 10.1101/2022.12.14.520403

**Authors:** Michał Brouze, Marcin Szpila, Areta Czerwińska, Wiktor Antczak, Seweryn Mroczek, Tomasz M. Kuliński, Anna Hojka-Osińska, Dominik Cysewski, Dorota Adamska, Jakub Gruchota, Ewa Borsuk, Andrzej Dziembowski

## Abstract

Among numerous enzymes involved in RNA decay, processive exoribonucleases are the most prominent group responsible for the degradation of the entire RNA molecules. The role of mammalian cytoplasmic 3’-5’ exonuclease DIS3L at the organismal level remained unknown. Herein we established knock-in and knock-out mouse models to study DIS3L functions in mice. DIS3L is indeed a subunit of the cytoplasmic exosome complex, which disruption leads to severe embryo degeneration and death in mice soon after implantation. These changes could not be prevented by supplementing extraembryonic tissue with functional DIS3L through the construction of chimeric embryos. Preimplantation *Dis3l*^-/-^ embryos were unaffected in their morphology and ability to produce functional embryonic stem cells showing that DIS3L is not essential for cell viability. There were also no major changes in the transcriptome level for both embryonic stem cells and blastocysts, as revealed by RNA sequencing experiments. Notably, however, DIS3L knock-out led to inhibition of the global protein synthesis. These results point to the essential role of DIS3L in mRNA quality control pathways crucial for proper protein synthesis during embryo development.

## Introduction

In recent years RNA decay pathways emerged as the RNA control mechanisms of importance comparable to transcription regulation or posttranscriptional modifications. While RNA degrading enzymes consist of different types and families, processive exoribonucleases are the most prominent group, as they are responsible for the degradation of entire transcripts during mRNA turnover or in quality control pathways. The mammalian genomes encode six processive exoribonucleases, divided into three families. The first family of 5’ to 3’ exonucleases consists of XRN1 and XRN2, expressed, respectively, in P-bodies scattered across cytoplasm (1) and nucleus (including nucleolus) (2). The second family consists of three hydrolitic 3’ to 5’ exoribonucleases. Two of them, nuclear DIS3 (which also retained endoribonucleolytic activity) and cytoplasmic DIS3L, are parts of the ring-shaped exosome complex (3,4), which by itself remains inactive in eukaryotes. Additionally, DIS3L2 exoribonuclease is also expressed in the cytoplasm, albeit in free, monomeric form (5). Finally, phosphorolytic 3’ to 5’ ribonuclease PNPase is responsible for RNA decay in mitochondria (6). While enzymes mentioned above are responsible for the bulk of RNA decay, plenty of less potent enzymes contribute to RNA metabolism, including nucleolar enriched, exosome associated distributive exoribonuclease EXOSC10/RRP6 (7,8). The substrates and mechanisms of recognition appearing in concert with activatory complexes are quite well established for those enzymes thanks to years of biochemistry, structural biology, and functional studies using cell lines. However, the knowledge about their role *in vivo* is more fragmentary.

DIS3 family of proteins, specifically, were shown to be associated with cancer. Mutation of DIS3L2, a nuclease responsible for the degradation of ARE-containing transcripts (5) and oligouridylated ncRNAs in their quality control pathway (9), is the leading cause of the Perlman syndrome and, consequently, Wilms tumor (10-12). Exosome-associated DIS3, which in the nucleus targets pervasive transcription products (promoter upstream transcripts and enhancer RNAs), snoRNAs and premature termination products (13), is among most frequently mutated genes in multiple myeloma patients (14-16). Unlike DIS3L2, however, we showed that DIS3 is an essential gene in mice, as evidenced by the analysis of both knock-out and point catalytic mutations of the catalytic domain (unpublished). Interestingly, this is also the case for the second nuclear exosome-bound exoribonuclease, RRP6, which was recently identified to be essential for mouse embryo development past the morula stage (17). Notably, essentially nothing is known about the role of DIS3L at the organismal level.

Here we present the results of our study of mammalian DIS3L *in vivo* through the analysis of knock-out mutation phenotype of *Dis3l* gene in mice. We demonstrate that DIS3L is essential for mouse embryo development beyond day 6,5. Lethal phenotype in *Dis3l* knock-out (KO) embryos could not be rescued by supplementation of wild-type (WT) cells to extraembryonic tissue. Although preimplantation embryos lacking functional DIS3L were able to produce embryonic stem (ES) cells, albeit, with reduced efficiency, they also displayed accumulation of a number of transcripts, as shown by RNA sequencing. Curiously, the accumulation of some of those transcripts does not produce higher protein levels. Further examination showed overall impaired protein production in those KO preimplantation embryos.

## Materials and Methods

### Animals

Mice lines were generated using CRISPR/Cas9 method in C57BL/6/Tar x CBA/Tar mixed background mice, following the procedure described previously (18). The loss-of-function *Dis3l*^-/-^ mutation was generated by random insertion of 169 bp fragment through non-homologous end joining repair mechanism in exon 10 in 1259 nt position of cDNA sequence of *Dis3l* gene (sgRNA sequence: TCCAGGTTGCCGTTATTCA). Knock-in *Dis3l*^*GFP/GFP*^ mutation was introduced in the ORF of *Dis3l* gene at the 3’-end through homologous recombination by insertion of GFP coding single-stranded DNA donor (sequence in Supplementary Figure 1.) with 60 bp homology arms on both ends (sgRNA sequence: GACAAAGGTCTTTAATGACA). All animal experiments were approved by the Local Ethical Committee in Warsaw affiliated to the University of Warsaw, Faculty of Biology (approval numbers: 527/2013, 176/2016) and were performed according to Polish Law (Act number 266/15.01.2015).

**Figure 1.**
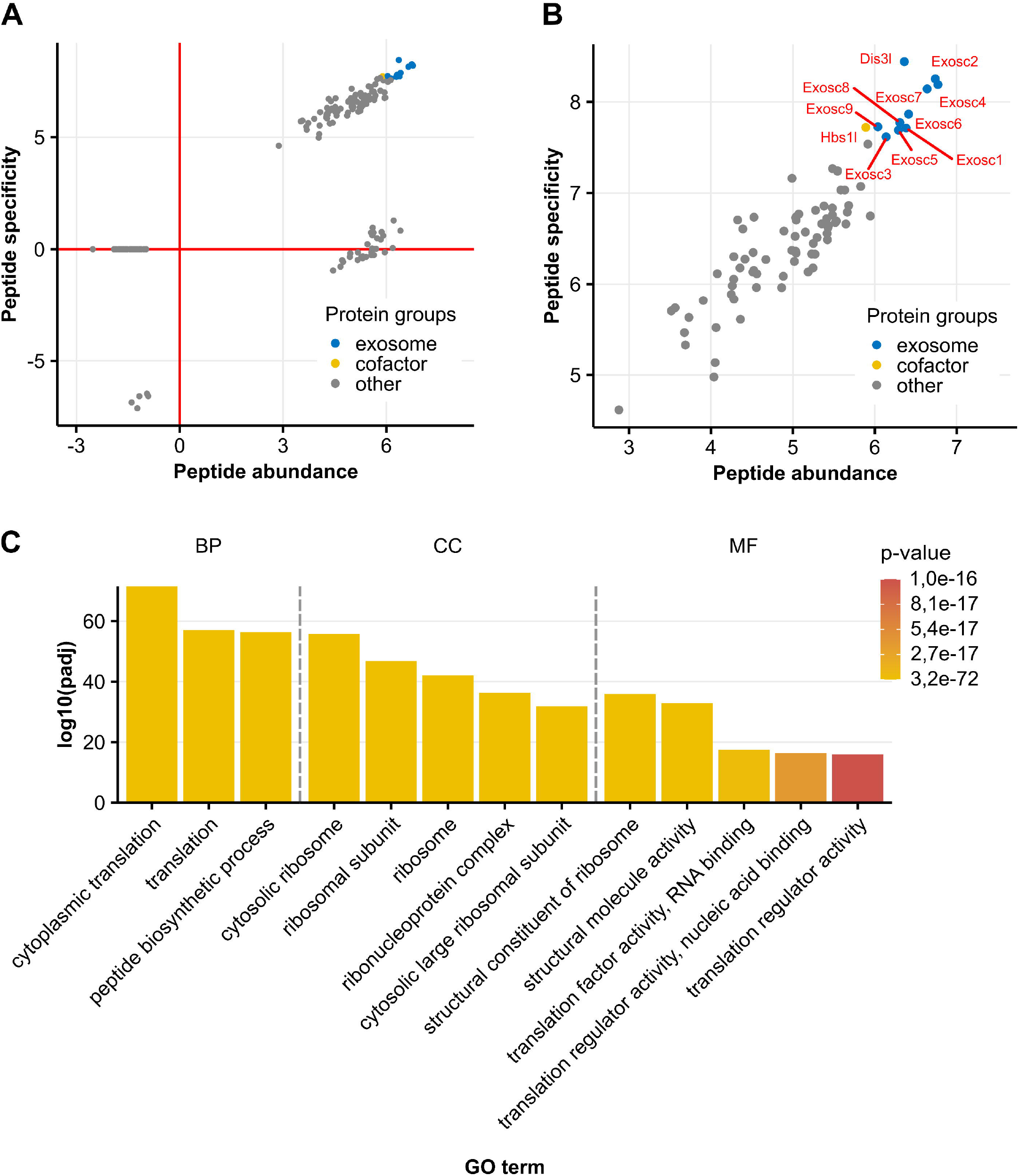
(**A**) Distribution of proteins identified by mass spectrometry analysis of DIS3L-GFP co-precipitated peptides from *Dis3l*^*GFP/GFP*^ protein liver extracts presented as peptide abundance relative to peptide specificity. (**B**) All nine mammalian core exosome proteins co-precipitated with DIS3L-GFP as the most specific and abundant proteins detected in the experiment. (**C**) Functional enrichment analysis of proteins co-precipitated with DIS3L-GFP. Only several terms with the highest p-value are presented. Identified terms are grouped by data source sub-groups: Biological Process (BP), Molecular Function (MF) and Cellular Component (CC). A full list of identified terms is available in Supplementary Figures 2-4.

### Embryos isolation and culture

Superovulation was artificially stimulated in females by intraperitoneal injection of 10 units of pregnant mare’s serum gonadotropin (PMSG, BioVendor) and human chorionic gonadotropin (hCG, MSD Animal Health), respectively, in 48 h interval. Stimulated females were mated with males in the evening and terminated by cervical dislocation and dissected in following days to obtain either 1-, 2-, 4-, 8-cell or blastocysts stage embryos. Zygotes were isolated by puncturing oviducts ampulla in hyaluronidase solution (Sigma-Aldrich) in PBS (300 μg/ml) to dissociate cumulus cells. Other embryos were isolated by flushing oviducts and uterus with M2 medium (Sigma-Aldrich). After isolation all embryos were rinsed in M2 medium and used for further procedures. Postimplantation embryos were isolated from unstimulated females 6 or 7 days after mating with males.

All embryos were cultured in cell culture incubator with 5% CO_2_ in the atmosphere and 38°C. Embryos were placed in drops of either M2 (for short-term culture) or M16 (for multi-day culture; Sigma-Aldrich) medium under mineral oil (Sigma-Aldrich) on 35-mm Falcon Easy-Grip tissue culture dishes (Thermo Fisher Scientific).

### Generation and in vitro culture of ES cells and EBs

For the derivation and culture of ES cells, feeder cells - inactivated mouse embryonic fibroblasts (MEFs) - were prepared according to Robertson (19). Blastocysts collected 96 h after hCG injection were transferred to single wells of culture plates coated with 0,2% gelatin (Sigma-Aldrich) with feeder layer of inactivated MEFs. Composition of the derivation medium was designed based on 2i method (20) (full composition in Supplementary Table 1). After 3-4 days of culture in cell culture incubator in 38°C and with 5% CO_2_ in the atmosphere, blastocysts formed outgrowths which were disaggregated by 5 min. incubation in 0,25% trypsin-EDTA (Gibco) followed by mechanical pipetting. Resulting cell suspensions were transferred into the separate culture plate wells onto fresh layer of inactivated MEFs and inspected daily for appearance of primary colonies. Cultures containing ES cells were expanded, processed for genotyping, and frozen for further investigation. Established ES cell lines were cultured in ES cells medium composed of KnockOut Dulbecco’s modified Eagle’s Medium (KnockOut DMEM, Gibco) supplemented with 15% heat inactivated FBS (Performance Plus, Gibco) with addition of nonessential amino acids, L-glutamine, β-mercaptoethanol, penicillin and streptomycin, and 500 IU/ml LIF (Gibco).

To gather pure ES cells population MEFs were removed from cultures by pre-plating, i.e. incubating cells with 0.05% trypsin-EDTA (Sigma-Aldrich) for 3-5 min and seeding of suspended cells onto culture dish covered with gelatin for 20 min, allowing MEFs to attach to the dish. MEF-free suspension of ES cells was cultured on a new gelatin-covered culture dish until reaching confluency. ES cells were then detached by incubation in 0,05% trypsin-EDTA for 3-5 min and washed in PBS (BioShop). Centrifugated dry pellets were frozen at -80°C until further processing.

To induce differentiation of ES cells, embryoid bodies (EBs) were generated. First, ES cells’ colonies were disaggregated using trypsin and pre-plated for 20 minutes (as described above). Part of collected cells suspension (day 0 of EB culture) was kept for RNA isolation by freezing a dry pellet after wash in PBS. The second part of cells suspension was centrifuged, and cells were cultured in differentiation medium (ES cells culture medium without LIF) in the concentration of 800 cells per 30 μl in hanging drops to stimulate EBs formation. After 2 days of culture in hanging drops EBs were either collected for analysis or washed and placed in low-adhesive dishes (Medlab) allowing for their culture in suspension. After 5 days of differentiation, the rest of EBs were collected for analysis.

### Immunoprecipitation

Livers from 10-20-week-old *Dis3l*^GFP/GFP^ knock-in mice were homogenized in lysis buffer (10 mM Tris-HCl pH 8; 125 mM NaCl; 1 mM DTT; 1 mM PMSF; 0,02 μM pepstatin A; 0,02 ug/ml chymostatin; 0,006 μM leupeptin; 20 uM benzamidine hydrochloride) and processed as described in (3) and (21) and analysed by mass spectrometry by the Mass Spectrometry Laboratory, IBB PAS. Identified proteins that were considered non-specific contaminants (haemoglobin subunits, immunoglobulins, keratins and trypsinogens) were excluded from the final analysis of enriched proteins.

### DNA isolation and genotyping

For genotyping of mice, postimplantation embryos and ES cells adjusted HotShot method was used (22). DNA from fixed and stained embryos was isolated using the protocol described in (23). Preimplantation embryos were repeatedly frozen and warmed several times in a minimal amount of RNase and DNase-free water in PCR tubes. Obtained DNA was used for genotyping by PCR reaction using Phusion High-Fidelity DNA polymerase (Thermo Fisher Scientific). Reactions of 20 μl volume were set up according to the manufacturer’s protocol, with no DMSO and 10 μM primers used (primer sequences in Supplementary Table 2). Additionally, blastocysts from which RNA was extracted for sequencing and RT-qPCR experiments, were genotyped as described for gDNA above, with cDNA generated during those experiments as a PCR template.

### RNA isolation and processing

RNA from ES cells was extracted by lysis of 1 million cells using TRI Reagent (Sigma-Aldrich), following manufacturer’s protocol. RNA from EBs was isolated using High Pure RNA Isolation Kit (Roche), following manufacturers protocol. RNA from living embryos was extracted using PicoPure RNA Isolation Kit (Thermo Fisher Scientific) following manufacturers protocol with following modifications: single embryos were incubated for 30 min at 42°C in Extraction Buffer and then mixed with ethanol in 1:1 ratio. All isolated RNA samples were stored at -80°C until further use. DNA was removed from all RNA samples using TURBO DNA-free Kit (Thermo Fisher Scientific) following manufacturers protocol for routine DNase treatment, with additional 2 μl of RiboLock RNase Inhibitor (Thermo Fisher Scientific) added to every reaction. For RNA sequencing and RT-qPCR experiments cDNA was synthesized. For single blastocysts RNA sequencing, RT-qPCR and ES cells RT-qPCR experiments, SuperScript III Reverse Transcriptase (Thermo Fisher Scientific) and oligo(dT)_20_ primer was used to generate cDNA from total RNA, following modified manufacturer’s protocol. Reverse transcription reactions were cleaned up using AMPure XP Reagent magnetic beads (Beckman Coulter). For EBs RT-qPCR, 250 ng of total RNA was used for reverse transcription reaction with RevertAid First-Strand cDNA Synthesis kit (Thermo Fisher Scientific), following manufacturer’s protocol.

### Chimeric embryos

To produce tetraploid (4n) WT embryos, 2-cell diploid (2n) WT embryos were isolated from superovulated females 30 h after hCG administration. Early 2-cells embryos were electroporated using home-made electroporation chamber and programmable square pulse generator. Embryos were rinsed several times and placed in electroporation buffer (100 mM MgSO_4_*7H_2_O, 100 mM CaCl_2_*2H_2_O and 0,045 g/ml glucose in H_2_O) warmed to 38°C in a glass petri dish between two straight electrodes of platinum wire attached to the dishes bottom approximately 100 μm apart, with embryo cleavage plane parallel to the electrodes. Two 40 V pulses lasting 40 μs in 100 μs interval were generated to induce fusion of the blastomeres. 2-cell embryos were washed once in M2 medium and placed in culture for 24 h. Within the first hour from electroporation embryos were inspected for survival and blastomere fusion and sorted adequately to trace their development back to 2-cell stage.

Simultaneously to 4n WT embryos reaching 2-cell stage, 4-to-8-cell 2n embryos were isolated from *Dis3l*^*+/-*^ females previously mated with males. All embryos had their zona pellucida removed by short (ca. 10 s) rinsing in Tyrode’s Acidic Solution (Sigma-Aldrich) warmed to 38°C. Two 4n and one 2n embryos were rinsed and then moved to fresh aggregation solution: 0,3mg/ml phytohemaglutynin (Sigma-Aldrich) in M2 medium without BSA (home-made). Embryos were manually attached together using mouth pipette. Chimeric embryos prepared this way were rinsed with M2 medium and cultured overnight in M16 medium. Chimeric embryos were then transferred to pseudopregnant females on the first day of pseudopregnancy, following the same procedure as for the mice line generation.

### Embryo fixation and staining

Preimplantation embryos were fixed in 4% PFA (Thermo Fisher Scientific) for 30 min and permeabilised in 0,5% Triton X-100 (Sigma-Aldrich) for 20 min. After fixation embryos were placed in drops of blocking solution (3% BSA (Santa Cruz Biotechnology) in PBST) on plastic dishes overnight before immunostaining or were stored that way until further procedures.

All immunofluorescence labelling was performed by incubating fixed embryos in blocking solution with appropriate primary antibody dilution (Supplementary Table 3) at 4°C overnight. Embryos were washed once in PBS and twice in blocking solution and incubated with secondary antibody (Supplementary Table 3) for 2 h at room temperature. After 30 min washing in PBS blastocysts were ready for imaging or additional staining. Methionine analogue labelling was performed by incubating embryos in 25 μM Click-iT AHA (L-Azidohomoalanine, Thermo Fisher Scientific) solution in M16 medium for 2 h and then fixing them as described above. AHA incorporated in proteins was detected using dedicated Click-iT Cell Reaction Buffer Kit (Thermo Fisher Scientific) and Alexa Fluor 594 Alkyne (Thermo Fisher Scientific) in 1 μM final concentration according to manufacturer’s protocol. After brief washing in 3% BSA and chromatin staining embryos were ready for imaging. Chromatin in embryos was stained with Hoechst 33343 (1:5000 dilution in PBS, 2 μg/ml, Sigma-Aldrich) for 15 minutes.

All stained embryos were imaged on uncoated 35 mm plastic dishes with glass bottom (MatTek) in drops of M2 medium using LSM 800 Confocal Laser Scanning microscope (Zeiss). Obtained images were analysed using ImageJ and Python software. Cell number in blastocysts was counted in confocal sections manually using ImageJ built-in tools. SLC37A2 and AHA intensities were gathered from maximum projection of individually scanned blastocysts using Python and ImageJ, respectively.

### RNA sequencing libraries

For ES cells’ RNA sequencing, total DNA-free RNA was ribodepleted using Ribo-Zero Gold rRNA-removal kit (Illumina). rRNA-free RNA samples were cleaned-up by precipitation with 3 M sodium acetate. Sequencing libraries were prepared using KAPA Stranded RNA-Seq Library Preparation Kit (KAPA Biosystems). Between each step samples were cleaned up using AMPure XP Reagent magnetic beads. Library quality and fragment size distribution were verified by electrophoresis in Agilent 2100 Bioanalyzer (Agilent Technologies Inc.).

RNA-seq libraries from single blastocysts’ DNA-free RNA samples were prepared by tagmentation reaction following published protocols (24,25) with various steps and amount of enzyme optimized for use of a home-made batch of Tn5 transposase produced in our laboratory. Briefly: Tn5 (0,25 mg/ml) was loaded with linker oliguncleotides Tn5ME-A/Tn5Me-rev and Tn5ME-B/Tn5Me-rev by mixing of 10 μl Tn5 with 0,5 μl of both linkers (0,35 μM) and incubating for 45 min in 23°C with shaking at 350 rpm. Right before the reaction setup, loaded Tn5 was diluted 10 times with nuclease-free water. 10 μl of freshly prepared tagmentation buffer (20 mM Tris-HCl pH 7,5; 20 mM MgCl_2_; 50% dimethylformamide (Sigma-Aldrich) were mixed with 5 μl of diluted Tn5 and 5 μl of cDNA, incubated for 3 min at 55°C in a preheated thermocycler and cooled to 10°C for 1 min. Reaction was inactivated by addition of 5 μl of 0,2% SDS (Sigma-Aldrich) and incubation for 5 min at room temperature (RT). Tagmented cDNA was purified with AMPureXP Reagent magnetic beads using 1:1,25 cDNA to beads ratio.

For library amplification KAPA HiFi HotStart ReadyMix PCR Kit (Roche) with addition of 5% DMSO was used. Number of cycles yielding the best results was determined experimentally and was further individually adjusted for each batch of prepared cDNA. Library quality and fragment size distribution were verified by electrophoresis using Agilent 2100 Bioanalyzer.

ES cells’ RNA libraries were sequenced on NextSeq 500 instrument (Illumina) in 2×75 cycles pair-end mode in Next Generation Sequencing Unit of the International Institute of Molecular and Cell Biology in Warsaw. Single blastocysts’ RNA libraries were sequenced on NovaSeq 6000 (Illumina) instrument in 2×100 cycles pair-end mode in Genomics Core Facility of Centre of New Technologies, University of Warsaw.

### RNA sequencing bioinformatic analysis

Biological triplicates were analysed for each genotype and reads were mapped to the reference mouse genome (GRCm38) using the STAR short read aligner (version STAR v2.7.10a) (26). Quality control, read processing and filtering, and visualization of the results were performed using custom scripts and elements of the RSeQC, BEDtools and SAMtools packages. Reads were counted to the Genecode v M6 basic annotation using featureCounts of the subread package (v2.0.1). Differential expression analyses were performed using the DESeq2 (version 1.34.0) Bioconductor R package (27).

Functional enrichment analysis was performed with g:Profiler using gprofiler2 package for R (database version: e106_eg53_p16_65fcd97) with following settings: g:SCS multiple testing correction; 0,05 significance threshold (28).

### RT-qPCR

RT-qPCR experiments on ES cells and blastocysts were performed using Platinum SYBR Green qPCR SuperMix-UDG (Invitrogen) in LightCycler 480 Instrument (Roche) according to the manufacturer’s protocol. All ES cells cDNA samples were diluted 10 times before using as a reaction template. All primers used are described in Supplementary Table 2. All biological samples were run in technical triplicates. Number of biological samples is denoted in the Results section.

RT-qPCR experiment on EBs was performed using TaqMan Gene Expression Master Mix (Life Technologies) in LightCycler 96 instrument (Roche), according to the manufacturer’s protocol. All RNA samples were diluted 4 times. All samples were run in technical duplicates. Specific TaqMan probes against 4 different transcripts were used: Mm00443081_m1 (*Pax6*), Mm01976556_s1 (*Foxa2*), Mm01318252_m1 (*Tbxt*), Mm01205647_g1 (*Actb*).

Cp values of all technical replicates of one biological sample results were averaged. These results were analysed using 2^-ddCp method (final formula used: 2^-ddCp = 2^-[(Cp_GOI_ – Cp_housekeeping_)_KO_ – (Cp_GOI_ – Cp_housekeeping_)_WT_]). Obtained Cp values of genes of interest from every sample were normalized to either actin b (EBs) or GAPDH (ES cells and blastocysts) Cp value. For blastocysts RT-qPCR analysis, only samples with technical replicates’ standard deviation below 1 were chosen.

### Statistical analysis

All statistical analyses were performed using RStudio. All data plots present individual data points (where applicable), as well as mean value with standard error of mean (SEM) indicated as error bars (where applicable). All experiments were performed in at least two individual repetitions (with exception of immunoprecipitation experiment with one repetition) and in each experiment biological samples were obtained from at least 3 different animals. Distribution ratios were compared by Fisher exact test for 2×2 contingency tables (viability of embryos) or Chi-square test for 2×3 tables (genotype distributions). Mean values were compared using either parametric two-sided t-test for sample sets with normal distribution or non-parametric two-sided Mann-Whitney-Wilcoxon test for sample sets without normal distribution. Distribution normality was tested with Shapiro-Wilk test. Value “n” in figures and figure descriptions denotes the number of individual embryos analysed. P-values for every test are reported on plots.

## Results

### DIS3L is a part of the cytoplasmic exosome in mice

The composition and functionality of the mammalian cytoplasmic exosome complex and its catalytic subunit, DIS3L, was previously analysed only in highly modified cell lines (HEK293 and HeLa). Thus, we have generated a knock-in mouse line with DIS3L C-terminally tagged with GFP in the endogenous locus using the CRISPR/Cas9 system to study it in a more physiologically relevant system. These mice in the homozygous state were healthy, with no apparent developmental, morphological, or physiological abnormalities. Then, we used protein extracts from three DIS3L^GFP/GFP^ mice livers, to determine DIS3Ls association with cytoplasmic exosome complex in mammals by performing immunoprecipitation on anti-GFP resin. Extracts from one WT mouse liver served as a control. Quantitative mass spectrometry analysis showed 90 proteins significantly enriched in the averaged results of three *Dis3l*^GFP/GFP^ samples. DIS3L co-precipitated with all nine core exosome proteins with the highest abundance as well as one of the cytoplasmic exosome’s cofactors – HBS1L (Figure 1A, B).

The majority of other proteins co-precipitating with the DIS3L are associated with the protein synthesis apparatus. This includes 32 ribosomal proteins and 9 of translation initiation factors of the EIF family as well as GYGIF2, a factor involved in translation quality control. This is also supported by the functional annotation analysis of identified proteins – Molecular Function, Biological Process and Cellular Component gene ontology terms identified with the highest p-value are all associated with the translation process and its regulation (Figure 1C; Supplementary Figures 2-4).

**Figure 2.**
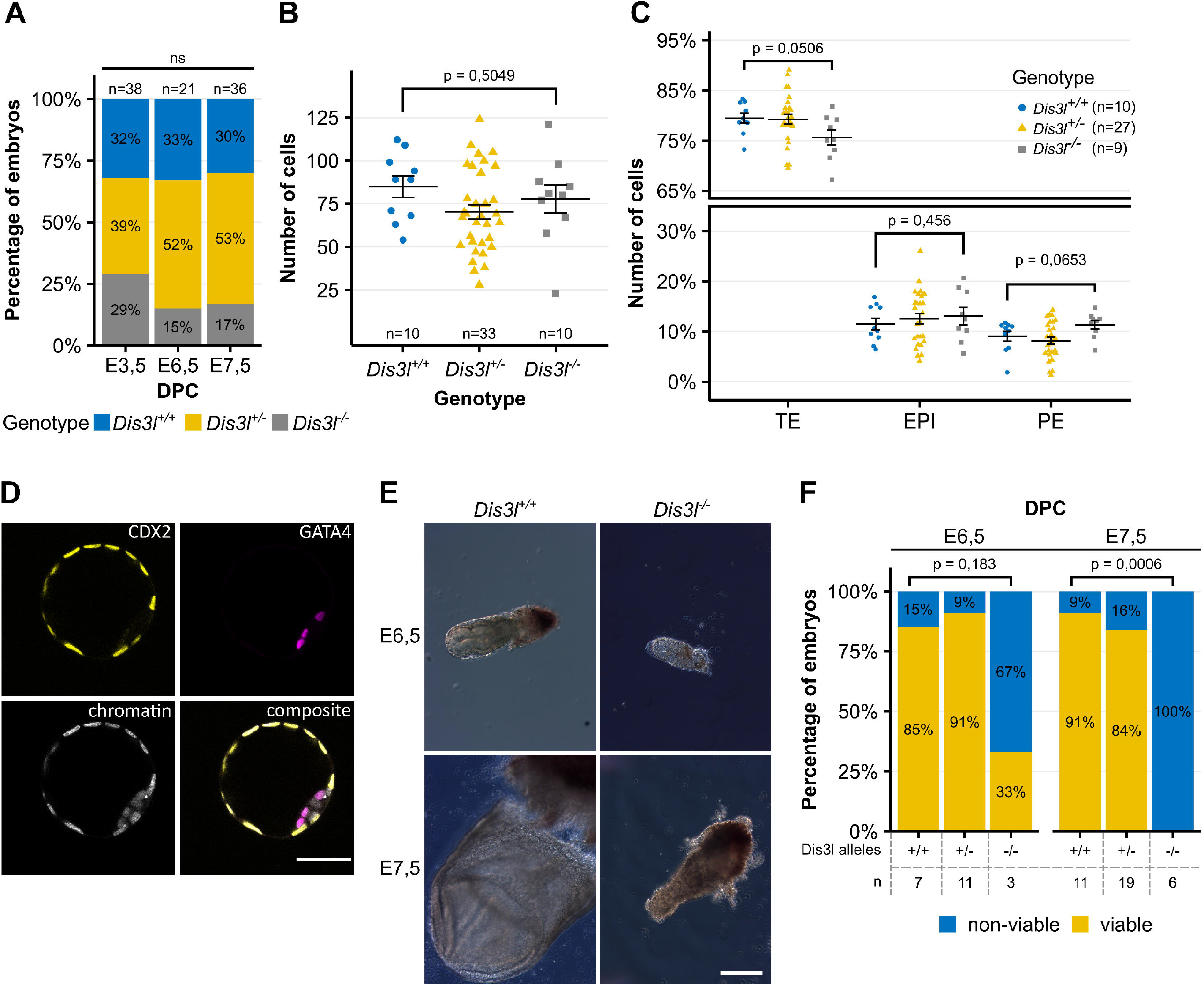
(**A**) Genotypes distribution in embryos of different developmental stages obtained from *Dis3l*^*+/-*^ x *Dis3l*^*+/-*^ matings. (**B**) Total cell count of blastocysts in relation to the genotype. (**C**) Percentage ratio of trophectoderm (TE), epiblast (EPI) and primitive endoderm (PE) in blastocysts in relation to the genotype. (**D**) Example of staining used to determine the number of differentiated cells in blastocyst. CDX2 marker of trophectoderm in yellow, GATA4 marker of primitive endoderm in magenta, chromatin in grey. Number of epiblast cells was determined by subtracting the number of CDX2 and GATA4 positive cells from total cell number. Scale bar = 50 um. (**E**) Example of incorrectly developing *Dis3l*^*-/-*^ embryos at E6,5 and E7,5 DPC compared to valid *Dis3l*^*+/+*^ embryos. Scale bar = 200 μm. (**F**) Ratio of viable to nonviable embryos of different genotypes at E6,5 and E7,5 DPC. Individual data points represent single blastocysts analysed with black summary bars representing mean value and error bars representing SEM.

Thus, DIS3L is indeed a subunit of the exosome complex, which in the cytoplasm is associated with the translation machinery.

### *Dis3l*^*-/-*^ embryos die at E6,5

Having established that DIS3L in mice is indeed a part of the cytoplasmic exosome complex, we aimed to determine its role in shaping mice transcriptome and its potential influence on mice physiology. Using CRISPR/Cas9 system, we generated a mice line bearing c.1258_1259ins169 frameshift mutation in the *Dis3l* gene, leading to disruption of the protein functionality. In our efforts to breed generated *Dis3l*^*-/-*^ mouse line, we were unable to obtain mice bearing homozygotic mutation. Therefore, we attributed it to embryolethality of homozygotic *Dis3l* mutation. Genotyping embryos at different developmental stages from breeding *Dis3l*^*+/-*^ males and females further confirmed the lethality.

Preimplantation development was unaffected in the progeny of those matings. The genotype ratio of obtained blastocysts did not differ significantly from the expected mendelian ratio, with 28,95% *Dis3l*^*-/-*^ and 31,58% *Dis3l*^*+/+*^ among them (Figure 2A). Lack of the functional cytoplasmic exosome also did not disturb the overall development of a preimplantation embryo. Both wild-type and KO blastocysts had similar mean total cell count (84,8 ± 6,27 and 77,8 ± 8,13 respectively, Figure 2B) and the ratio of all three cell lineages (i.e. trophectoderm, epiblast, and primitive endoderm) present at that stage of development (Figure 2C, D).

We observed the first signs of impaired development early after implantation. On days 6,5 and 7,5 of embryo development, we identified only 14,29% and 16,67% *Dis3l*^*-/-*^ embryos, respectively, although the entire genotype distribution on both stages remained within a range similar to the mendelian ratio (Figure 2A). However, the lack of functional DIS3L has already affected embryo morphology and developmental potential. While on day 6,5, more than 85% of *Dis3l*^*+/-*^ and *Dis3l*^*+/+*^ embryos were properly developed, 2 out of 3 identified KO embryos were smaller and deformed compared to their WT counterparts. This effect became stronger at day 7,5 when all 6 identified KO embryos were highly degenerated and non-viable for further development, without any change in the viability of embryos of other genotypes (Figure 2E, F).

It is concluded that the *Dis3l* KO mutation leads to embryo lethality at the implantation stage, but the cause of it remains to be established.

### Embryolethality of *Dis3l* KO cannot be rescued by supplementing WT extraembryonic tissues

One of the possibilities for the cause of *Dis3l* KO early embryolethality is the indispensability of *Dis3l* for the development of extraembryonic tissues responsible, among other functions, for proper implantation. To verify this, we constructed chimeric embryos. Individual 4-8-cell embryos (of unknown genotype) from *Dis3l*^*+/-*^ x *Dis3l*^*+/-*^ mating were joined with two 2-cell tetraploid WT embryos (Figure 3A). In such a model, tetraploid cells contribute only to the development of extraembryonic tissue – compensating for the lack of DIS3L in KO embryos and potentially rescuing the phenotype. Chimeric embryos were then transferred to recipient females previously mated with vasectomized males. All females were euthanised 18 days after an embryo transfer to account for all implanted embryos and resorption sites in the uterus.

**Figure 3.**
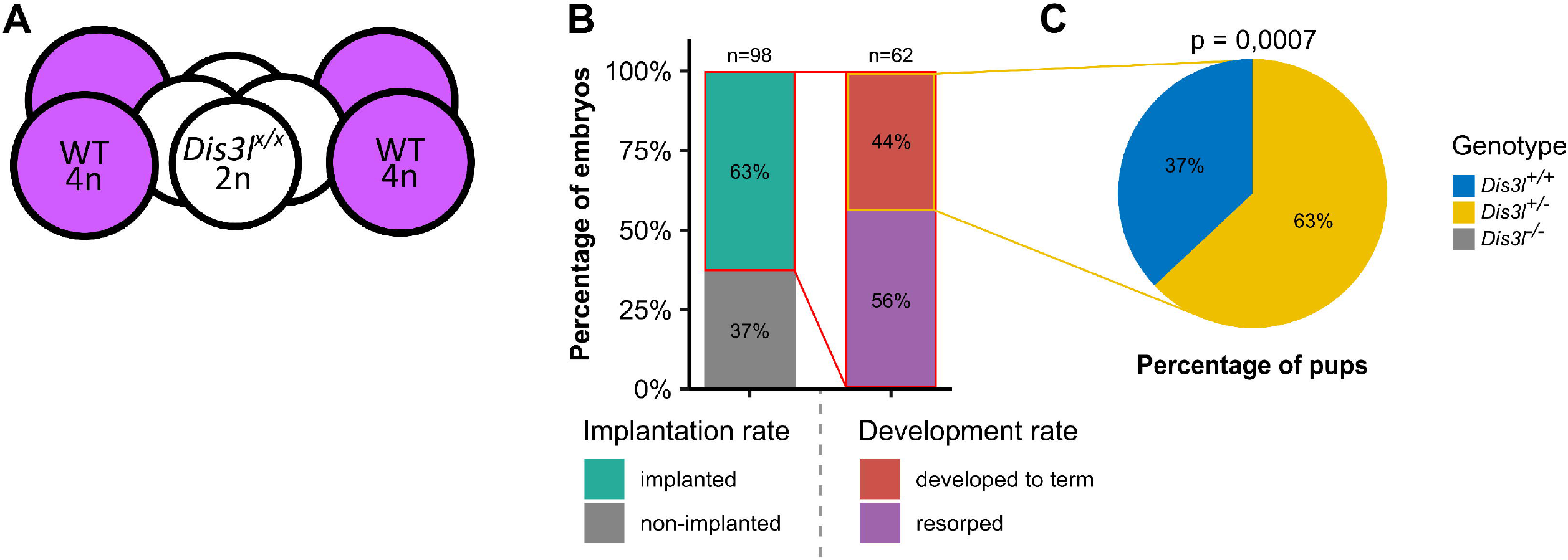
(**A**) Schematic representation of chimeric embryo construction. Diploid 4-cell embryos from *Dis3l*^*+/-*^ x *Dis3l*^*+/-*^ mating (white) were covered with two previously prepared tetraploid 2-cell wild-type embryos (magenta). (**B**) Chimeric embryos implantation rate after transfer to recipient females and percentage of those implanted embryos that fully developed to term. (**C**) Genotype distribution of chimeric pups developed to term. No *Dis3l*^*-/-*^ embryos developed fully.

Out of the total of 98 chimeric embryos transferred to recipient females, 63% of embryos were successfully implanted in the uterus. However, only 44% of those embryos developed to term, and the other 56% (35 out of 62) died and were subsequently resorbed, leaving only implantation sites (Figure 3B). Genotyping of developed embryos revealed that our attempt did not produce any *Dis3l*^*-/-*^ embryos, resulting in 67% of *Dis3l*^*+/-*^ and 33% of *Dis3l*^*+/+*^ pups developed to term(Figure 3C), all of them without any morphological signs of disturbed development. These results exclude the notion that the implantation process and, more broadly, the development of trophectoderm is the only cause of embryolethality.

### *Dis3l* KO does not affect cell viability

Having established that *Dis3l*^*-/-*^ embryo cannot develop properly after implantation and degenerates soon after, even with DIS3L supplemented in extraembryonic tissues, we assumed that it may be related to changes at the transcriptome level at this developmental stage. Thus, we next focused on analysing the effect of *Dis3l* KO mutation on the developmental potential of the preimplantation embryo.

To test the potential of the inner cell mass (ICM) of the preimplantation embryo to develop the embryo body, we first derived ES cell lines from blastocysts obtained from *Dis3l*^*+/-*^ x *Dis3*^*+/-*^ matings. We observed that *Dis3l*^*-/-*^ ES lines are obtained with a slightly lower ratio compared to *Dis3l*^*+/-*^ and *Dis3l*^*+/+*^ embryos. Of 28 blastocysts from which we derived ES cells, 14,29% were *Dis3l*^*-/-*^ (Figure 4A). Notably, both KO and WT ES cells were also able to form embryoid bodies. However, out of three tested KO cell lines, two produced embryoid bodies that, on days 2 and 5 of culture, displayed a decrease in expression of *Foxa2* and *Tbxt* transcripts coding proteins responsible for differentiation of endodermal and mesodermal organs, respectively. In addition, all three lines also displayed an atypical expression pattern of Pax6, involved in ectoderm development, with a spike of transcript accumulation on day 2 of culture and a drop below starting level on day 5 (Figure 4B).

**Figure 4.**
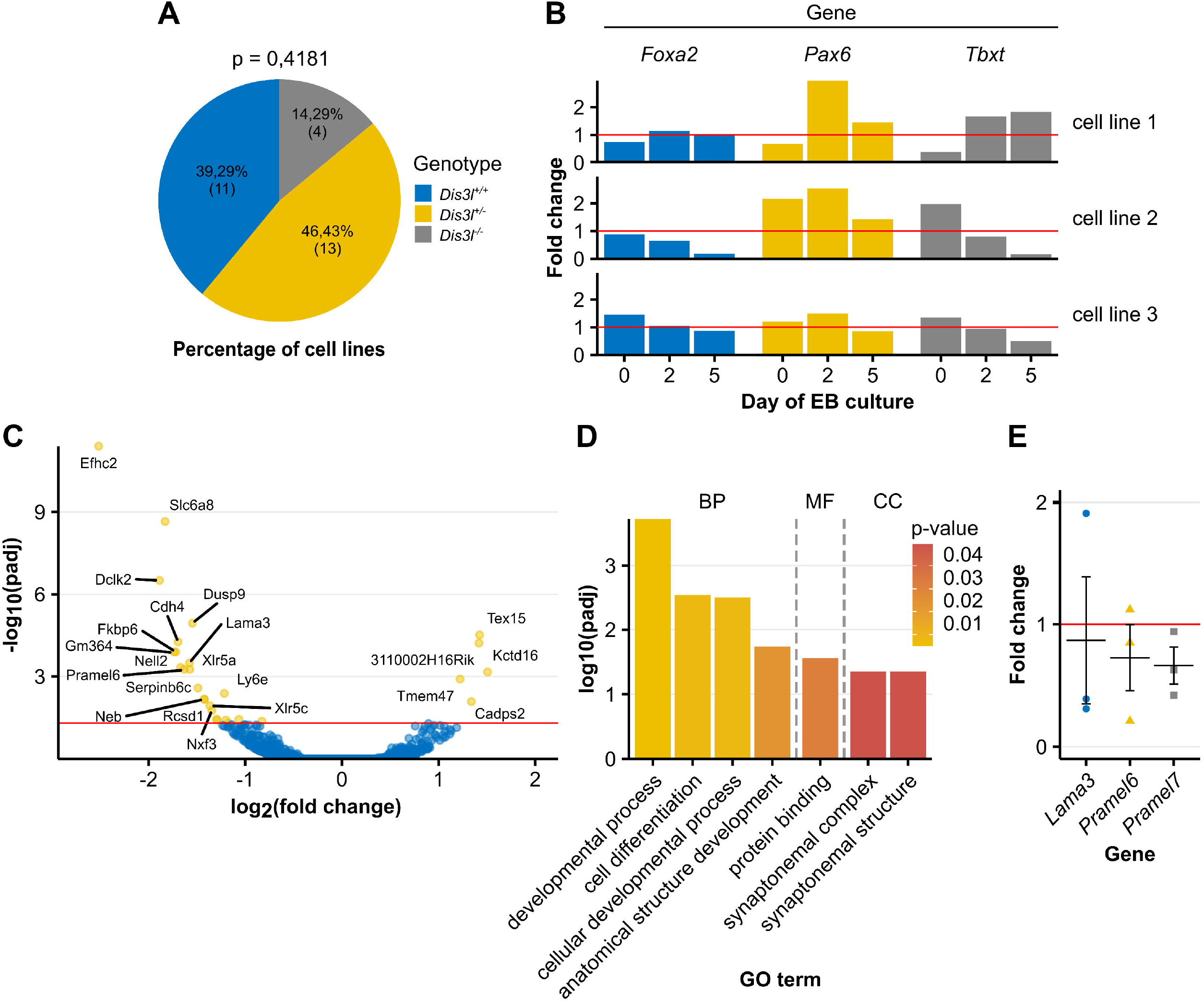
(**A**) Genotype distribution of ES cell lines derived from blastocysts from *Dis3l*^*+/-*^ x *Dis3l*^*+/-*^ matings. (**B**) Transcript level fold change of three germ layer-specific differentiation factors: *Foxa2, Pax6* and *Tbxt*, in embryoid bodies through 5 days of culture in three obtained *Dis3l*^*-/-*^ ES cell lines related to mean result of three WT cell lines, obtained by RT-qPCR experiment. Red horizontal line marks fold change of 1, meaning no change compared to WT. (**C**) Differential expression analysis of RNA sequencing results of three *Dis3l*^*-/-*^ and three *Dis3l*^*+/+*^ ES cell lines. Positive and negative log_2_(fold change) value represents up- and downregulation, respectively, of transcript in KO cells. Red horizontal line marks adjusted p-value of 0,05 with transcripts above it considered significantly changed in KO cells compared to WT ones. (**D**) Functional enrichment analysis of significantly deregulated transcripts identified in ES cells’ RNA sequencing. Identified terms are grouped by data source sub-groups: Biological Process (BP), Molecular Function (MF) and Cellular Component (CC). (**E**) Fold change of three of significantly downregulated transcripts identified in ES cells’ RNA sequencing in *Dis3l*^*-/-*^ ES cells related to mean result of three WT cell lines, obtained by RT-qPCR. Each data point represents fold change for one KO cell lines. Horizontal red line marks fold change of 1, meaning no change. Summary black bar represents mean fold change of a given transcript and error bars represent SEM.

To assess the state and possible changes in the transcriptome of ES cells lacking functional cytoplasmic exosome, we have performed a total RNA sequencing experiment on the Illumina platform, using RNA isolated from initial cultures. Comparing three *Dis3l*^*-/-*^ and three *Dis3l*^*+/+*^ ES cells lines, we identified 28 differentially expressed transcripts (out of all 13009 identified). 23 were downregulated and 5 upregulated (differential expression analysis, padj < 0,05) in KO cells. Of those, 20 had at least 2-fold expression difference (|log_2_foldchange| > 1) (Figure 4C). Gene ontology analysis revealed developmental and differentiation processes as the most enriched terms among those deregulated transcripts (Figure 4D). Among those downregulated, 3 genes (*Pramel6, Pramel7* and *Lama3*) are known to play a significant role in the regulation of ES cells and embryo development. *Pramel6* and *Pramel7* are essential for maintaining pluripotency of ES cells (29) and *Lama3* is a component of basal membrane during postimplantation development although it was only identified in decidua and Reichert’s membrane (30). To confirm their downregulation in KO samples, we compared the level of their transcripts in three WT and three KO samples by RT-qPCR. High variability of results in each biological sample was observed. For every gene tested, some samples showed upregulation, while others – downregulation of a given transcript (Figure 4E). This may be due to the process of differentiation being initiated at the time of cell collection.

It is concluded that although it is possible to derive and maintain a culture of *Dis3l* KO ES cells and those cells have very subtle changes on the transcriptome level, they display some abnormalities affecting differentiation factors which are difficult to explain mechanistically.

### *Dis3l* KO mutation negatively affects global protein production in blastocysts

Since the analysis of ES cells did not reveal the cause of the lethality-inducing effect of *Dis3l* KO mutation, we turned back to embryos at the blastocyst stage. Using a modified single-cell RNA sequencing protocol, we sequenced RNA of three *Dis3l*^*-/-*^ and three *Dis3l*^*+/+*^ blastocysts. Similarly to ES cells sequencing, we identified only a small number of differentially expressed transcripts (15 out of 17368 all identified). This time, however, only 2 of them were downregulated, and 13 – were upregulated (differential expression analysis, padj < 0,05). All these transcripts were also changed at least 2-fold (|log_2_foldchange| > 1) (Figure 5A), and the upregulated ones may represent direct substrates of the cytoplasmic exosome. Out of four Biological Process terms that identified deregulated transcripts were annotated by gene ontology analysis to, three regarded cell death (the same seven transcripts annotated to each). Additionally, three transcripts were successfully annotated to five different Human Phenotype terms (one transcript annotated to five terms, one to four and one to three), out of which four are related to developmental disorders (Figure 5B).

**Figure 5.**
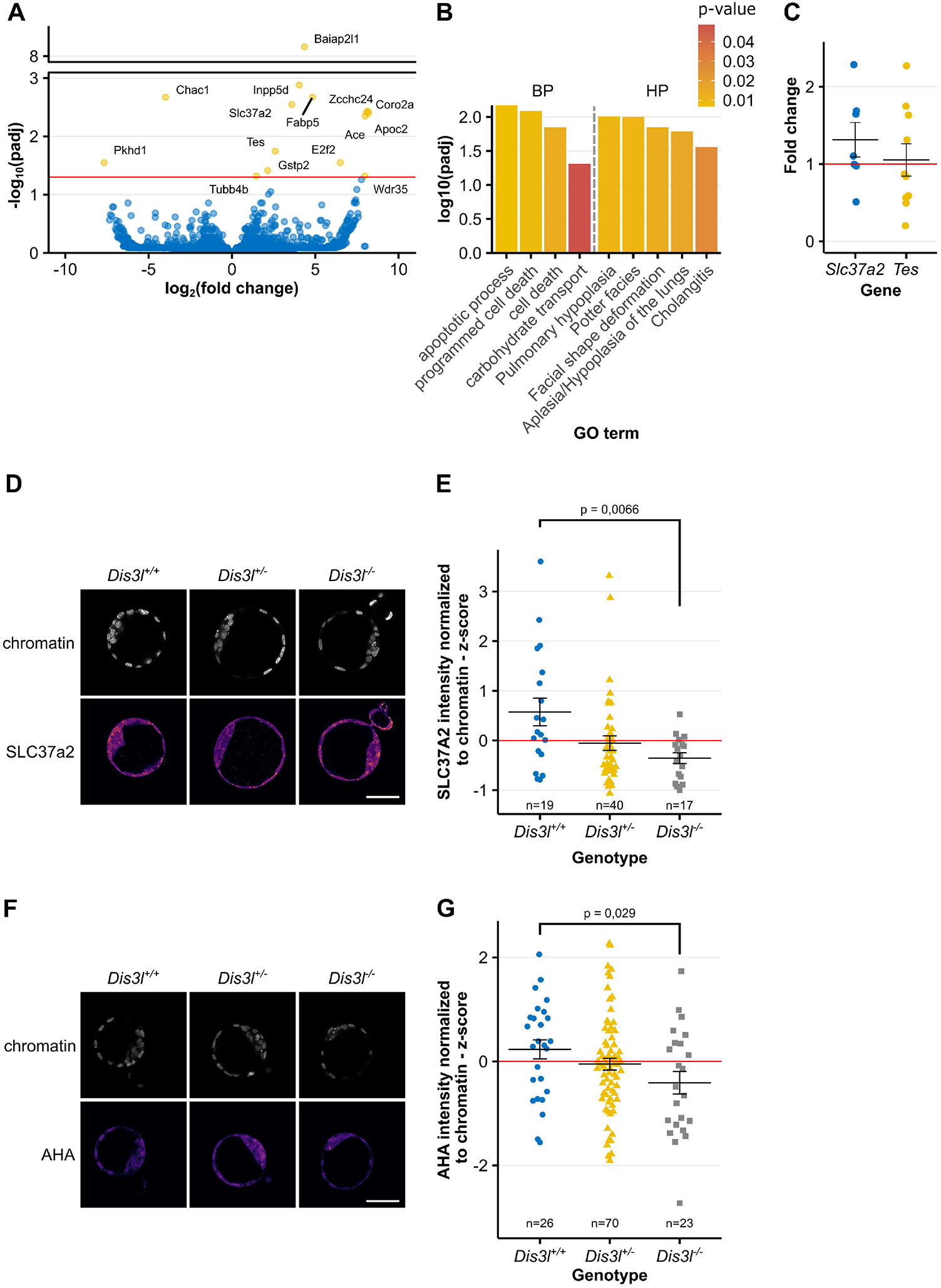
(**A**) Differential expression analysis of RNA sequencing results of three *Dis3l*^*-/-*^ and three *Dis3l*^*+/+*^ blastocysts. Positive and negative log_2_(fold change) value means up- and downregulation, respectively, of transcript in KO embryos. Red horizontal line marks adjusted p-value of 0,05 with transcripts above it considered significantly changed in KO embryos compared to WT ones. (**B**) Functional enrichment analysis of significantly deregulated transcripts identified in blastocysts RNA sequencing. Identified terms are grouped by data source sub-groups: Biological Process (BP) and Human Phenotype (HP). (**C**) Fold change of two of significantly downregulated transcripts identified in blastocyst’s RNA sequencing of *Dis3l*^*-/-*^ embryos, obtained by RT-qPCR. Each data point represents fold change for one KO blastocyst. Horizontal red line marks fold change of 1, meaning no change. (**D**) Example of SLC37A2 protein immunofluorescence staining used to measure protein level in blastocysts. (**E**) Normalized z-score value of SLC37A2 protein level in blastocysts depending on genotype. Each data point represents one blastocyst. (**F**) Example of protein synthesis labelling with methionine analogue AHA used to determine the amount of newly produced peptides in blastocysts. (**G**) Normalized z-score value of detected AHA-containing proteins levels in blastocysts depending on genotype. Each data point represents one blastocyst. Scale bars = 50 μm. Black summary bars represent mean value and error bars represent SEM.

Notably, two genes upregulated in *Dis3l*^*-/-*^ blastocysts are directly involved in preimplantation and early postimplantation development. Testin (*Tes*) is known for interacting with the cytoskeleton and affects actin dynamics to facilitate cell movements and migration (31,32). In blastocysts, it’s highly expressed in ICM and polar trophectoderm and after implantation at E6,5 in epiblast, anterior visceral endoderm precursors and extraembryonic ectoderm (33). SLC37A2 is a glucose-6-phosphate antiporter responsible for its transport from cytoplasm to ER lumen (34) – one of the steps in the metabolism of the primary blastocyst energy source – glucose (35,36). RT-qPCR analysis of those two transcripts’ levels from seven *Dis3l*^*-/-*^ and ten *Dis3l*^*+/+*^ blastocysts revealed that while *Slc37a2* transcript was accumulated in most of KO samples, with mean fold change of 1,31 ± 0,22, mean fold change of Testin-coding transcripts was only 1,05 ± 0,2 (Figure 5C). Thus, we focused on *Slc37a2*. Surprisingly immunofluorescence staining of SLC37A2 protein in blastocysts revealed that an elevated amount of transcript did not produce higher protein levels. On the contrary, standardized z-score of SLC37A2 protein amount in *Dis3l* KO blastocysts significantly decreased compared to WT embryos (mean z-score -0,35 ± 0,10 and 0,57 ± 0,28, respectively) (Figure 5D, E). All this suggests that the lethality of *Dis3l* KO is not related to changes in the transcript levels.

Decreased levels of SLC37A2 protein despite increased mRNA expression and the fact that cytoplasmic exosome is connected with the cytoplasmic quality control pathways may suggest global problems with protein synthesis resulting from stress responses. To measure the global protein synthesis levels, we performed metabolic labelling with clickable methionine analogue L-Azidohomoalanine (AHA). Quantification of AHA incorporation in newly synthesised proteins in embryos revealed the negative impact of *Dis3l* KO mutation on global protein synthesis in the preimplantation embryo (mean z-score -0,41 ± 0,22 and 0,23 ± 0,18 for WT embryos; Figure 5F, G).

It is concluded that *Dis3l* KO leads to the inhibition of protein synthesis in early embryos with a substantial effect on mRNA levels

## Discussion

In this paper, we studied the role of mammalian DIS3L processive exoribonuclease at the organismal level. DIS3L, which co-precipitates with all nine core exosome proteins (8), is essential for mouse embryo development. *Dis3l* KO mutation causes severe embryo degeneration and death between day 6,5 and 7,5 of its development. This is due not only to the implantation process mediated by extraembryonic tissue, but mostly to the changes in subcellular processes that occur before implantation. While KO preimplantation embryos do not display any changes in morphology or differentiation of first embryonic cell lineages and ability to produce ES cells from ICM, the translation rate was reduced, despite small changes at the transcriptome level. This presents a picture of the highly deregulated state of the early embryo at the level of subcellular processes.

Despite the disruption of global protein production, *Dis3l*^*-/-*^ embryos are still able to implant in the uterus and to function there for a brief period, but their fate is most certainly determined at the preimplantation stage. The state of both relies heavily on the mRNA and translation quality control pathways in the cell, in many of which DIS3L, as a catalytic subunit of cytoplasmic RNA exosome complex, is intrinsically involved (most notably no-go (NGD), nonsense-mediated (NMD) and non-stop (NSD) decay pathways of RNA degradation (37-39)). Thus, the upregulation of transcripts observed by us in blastocysts with mutated *Dis3l* would rather stem not from the elevated expression of certain genes but rather from the accumulation of aberrant transcripts that the functional exosome would normally degrade. As for proteome, translational machinery could still be recruited to those lingering transcripts but produced protein products would be incomplete or incorrect and, in return, would also have to be degraded by translation control mechanisms. Many of those mechanisms are triggered by ribosome stalling and collision events on abnormal mRNA particles. In 2020, two groups independently showed that SKIV2L in Ski complex, responsible for unwinding and feeding RNA molecules into the exosome channel (40-42) is required for the recruitment of the exosome in mRNA quality control pathways through its interaction with stalled ribosomes. This allows for their extraction through degradation of stalling-causing mRNA particles, making those ribosomes available for dissociation and recycling by Pelota/Hbs1/ABCE1 complex (43). Hence, the debilitating effect of the cell’s inability to degrade those problematic mRNAs would be two-fold: occurring ribosome collision events could not be properly resolved, and the number of stalling-causing mRNA particles would grow in time, causing new collision events.

While local mechanisms recruited to colliding ribosomes counteract negative consequences of these events, like EDF1 and GIGYF2 working together to inhibit translation from mRNA particles on which collision occurred (44,45), they can be overloaded by the global scale of the occurring problems. In such a case, more general stress response mechanisms are triggered, leading to global translation initiation block by the action of GCN2 and eIF2α phosphorylation or, in a case of stress being unresolved by the cell, apoptosis activated by MAPKKK ZAKα (46). As we present, this is the case for *Dis3l*^*-/-*^ preimplantation embryos, where the global protein amount is lowered despite the accumulation of multiple transcripts. Additionally, recently SKI2VL was also shown to be essential for mouse development, with whole-body deletion leading to embryolethality before day E13,5 (47). Thus, we propose that DIS3L, while dispensable for cell viability, is essential for embryo development as a part of RNA quality control response for translational stress, with its mutation possibly causing translation inhibition across the preimplantation embryo, and, further on, due to accumulation of the stress-inducing events, wide apoptosis after the implantation, leading to embryo death after day E6,5.

## Data Availability

The mass spectrometry proteomics data have been deposited to the ProteomeXchange Consotrium via the PRIDE (48) partner repository with the dataset identifier PXD038745.

RNA sequencing data been deposited in NCBI’s Gene Expression Omnibus (49) and are accessible through GEO Series accession number GSE220800.

Other numerical data underlying results presented are available in the article and its online supplementary materials.

## Funding

This work was supported by National Science Centre (grant numbers UMO-2013/10/M/NZ4/00299; UMO-2016/22/A/NZ4/00380), Foundation for Polish Science (grunt number TEAM TECH CORE FACILITY/2017-4/5) and European Union’s Horizon 2020 research and innovation program (grant agreement no. 810425).

## Conflict of Interest

Authors report no conflict of interest.

## Author Contribution

M.B. performed embryo genotyping, blastocyst and ES cells qPCR experiments and prepared RNA-sequencing libraries, M.B. and M.S. constructed and transferred chimeric embryos, A.C. derived ES cells and performed EB experiments, M.B and W.A. performed SLC37A2 and AHA labelling experiments, S.M. performed IP experiment, A.H.-O. and T.K. performed differential expression analyses, D.C. performed and analysed mass spectrometry, D.A. performed sequencing experiments, J.G. and E.B. produced all mice lines, M.B. and A.D. wrote the manuscript, A.D. conceived the study.

## Acknowledgments

We thank Andrzej Dziembowski lab members for their support and fruitful discussions, Anna Ciemerych for the support in the experiments involving ES cells and postimplantation embryos, Olga Gewartowska for multiple suggestions and critical reading of the manuscript, Aleksandra Brouze for critical and proofreading of the manuscript and all members of Genome Engineering Unit of International Institute of Molecular and Cell Biology in Warsaw for maintenance of animal colony and animal genotyping. NGS was performed thanks to Genomics Core Facility CeNT UW (RRID:SCR_022718), using NovaSeq 6000 platform financed by Polish Ministry of Science and Higher Education (decision no. 6817/IA/SP/2018 of 2018-04-10).

## Notes

### Competing Interest Statement

The authors have declared no competing interest.

