## Supplementary Materials for "DIS3L, cytoplasmic exosome catalytic subunit, is essential for development but not cell viability in mice"

**Supplementary Table 1.**

| Component | Ratio/concentration | Unit | Manufacturer |
|---|---|---|---|
| Neurobasal | 2 | - | Gibco |
| DMEM | 1 | - | |
| F-12 | 1 | - | |
| N2 | - | - | |
| B27 | - | - | |
| L-glutamine | 2 | mM | |
| Non-essential amino acids | 0,1 | mM | |
| BSA (Bovine Serum Albumine) | 0,1 | % | Sigma-Aldrich |
| β-mercaptoethanol | 0,1 | mM | |
| penicilin | 5000 | Units/ml | Gibco |
| streptomycin | 5000 | Units/ml | |
| LIF (Leukemia Inhibitory Factor) | 1000 | IU/ml | Chemicon International |
| MEK1 inhibitor - PD0325901 | 1 | uM | Sigma-Aldrich |
| GSK3 inhibitor – CHIR99021 | 3 | uM | Tocris Bioscience |

**Supplementary Table 1.** Medium composition for ES cells derivation. Components above thick bar are media composing final medium used, mixed in ratio denoted in Ratio/concentration column. Components below thick bar are additional components supplemented to the medium.

**Supplementary Table 2.**

| Target | Application | Direction | 5' -> 3' sequence |
|---|---|---|---|
| Dis3l | gDNA genotyping | forward | CGATGCTTGGGTTTCTGAAT |
| Dis3l | gDNA genotyping | reverse | TTGGAGGAAACAAACGATGG |
| Dis3l | cDNA genotyping | forward | AGCCCAGATGTGTGAGATGC |
| Dis3l | cDNA genotyping | reverse | AAGTGTGTGACGTCAGCGAT |
| Gapdh | RT-qPCR | forward | AAGGGCTCATGACCACAGTC |
| Gapdh | RT-qPCR | reverse | GGATGACCTTGCCCACAG |
| Slc37a2 | RT-qPCR | forward | GAAGGGAAAGCGGGGATTCA |
| Slc37a2 | RT-qPCR | reverse | ATGAATGACAGGCCCCAGTG |
| Tes | RT-qPCR | forward | AAGTACACCACCCTGATCGC |
| Tes | RT-qPCR | reverse | GGGGAGCCCACTCATAGGTA |

**Supplementary Table 2.** Primer sequences used in the study.

**Supplementary Table 3.**

| Staining | Antigen | Source | Producer | Cat. Number: | Dilution | Fluorochrome |
|---|---|---|---|---|---|---|
| Gata4 | Gata4 | goat | R&D Systems | AF2606 | 1:200 | - |
| Cdx2 | Cdx2 | rabbit | Abcam | ab76541 | 1:5000 | - |
| Slc37a3 | Slc37a2 | rabbit | Abcam | ab223048 | 1:200 | - |
| Gata4 | Goat IgG | donkey | Invitrogen | A-11055 | 1:500 | AF488 |
| Cdx2 | Rabbit IgG | donkey | Invitrogen | A-31573 | 1:500 | AF647 |
| Slc37a2 | Rabbit IgG | donkey | Invitrogen | A-10042 | 1:500 | AF568 |

**Supplementary Table 3.** Antibodies used in the study. Only secondary antibodies were conjugated with fluorophore.

**Supplementary Figure 1.**

ACACTTCTGGAGGAGATAAGGGACCTAGCTCTTCTGGATGTCTCTGACAGTTGTGCAATGgagaatttgtattttcag
ggtgatatcatggtgagcaagggcgaggagctgttcaccggggtggtgcccatcctggtcgagctggacggcgacgtaaacggccacaagttc
agcgtgtccggcgagggcgagggcgatgccacctacggcaagctgaccctgaagttcatctgcaccaccggcaagctgcccgtgccctggccca
ccctcgtgaccaccctgacctacggcgtgcagtgcttcagccgctaccccgaccacatgaagcagcacgacttcttcaagtccgccatgcccgaa
ggctacgtccaggagcgcaccatcttcttcaaggacgacggcaactacaagacccgcgccgaggtgaagttcgagggcgacaccctggtgaac
cgcatcgagctgaagggcatcgacttcaaggaggacggcaacatcctggggcacaagctggagtacaactacaacagccacaacgtctatatc
atggccgacaagcagaagaacggcatcaaggtgaacttcaagatccgccacaacatcgaggacggcagcgtgcagctcgccgaccactacca
gcagaacacccccatcggcgacggccccgtgctgctgcccgacaaccactacctgagcacccagtccgccctgagcaaagaccccaacgagaa
gcgcgatcacatggtcctgctggagttcgtgaccgccgccgggatcactctcggcatggacgagctgtacaagtaaTGAAATACTTCCATG
TCATTAAAGACCTTTGTCTTAAGTGGTGTACTTTTTTTTCTTTCT

**Supplementary Figure 1.** GFP donor sequence with homology arms (in uppercase).

**Supplementary Figure 2**

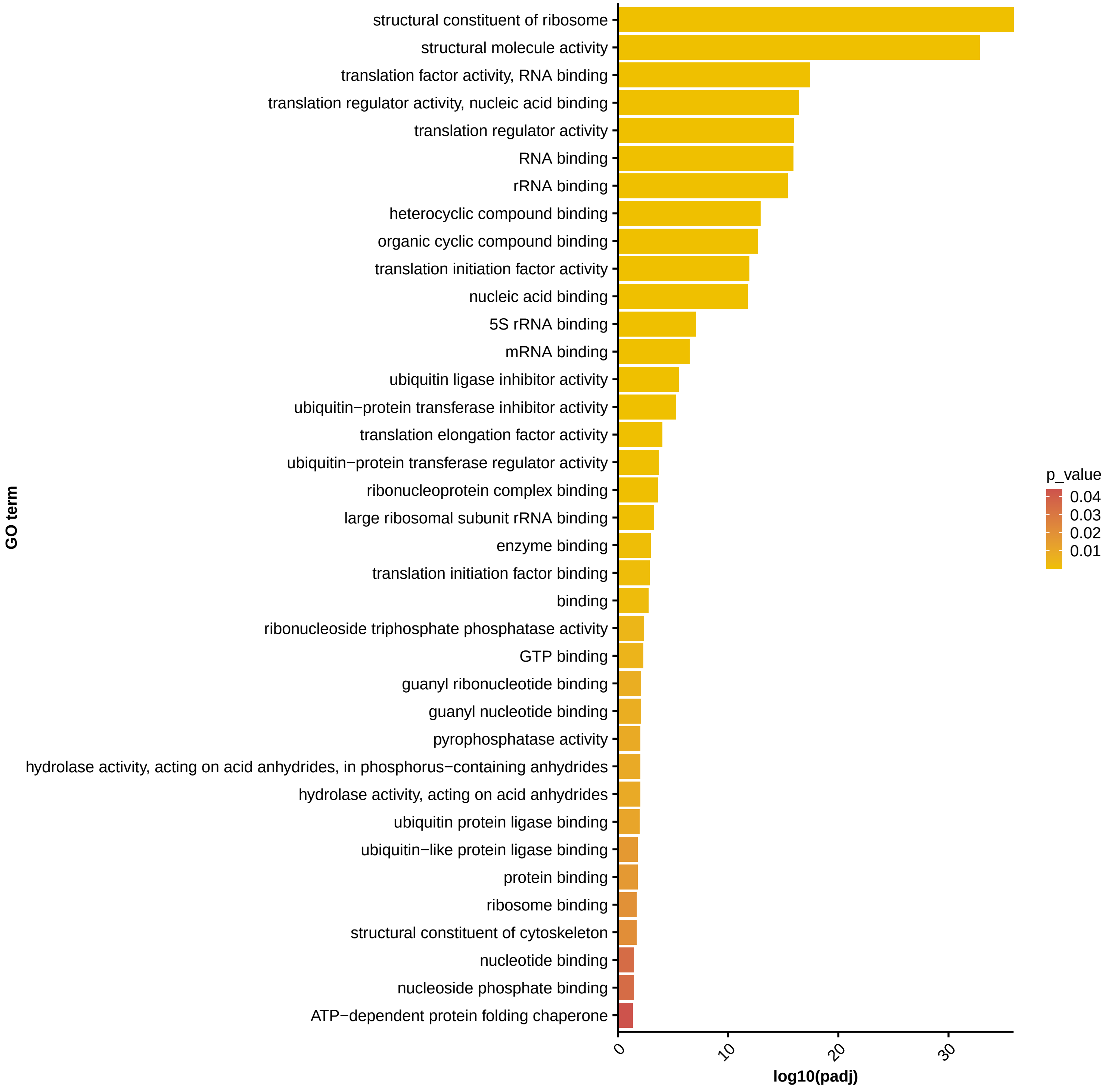

**Supplementary Figure 2.** Gene Ontology Molecular Function terms identified in functional annotation analysis of proteins coprecipitated with DIS3L-GFP.

**Supplementary Figure 3**

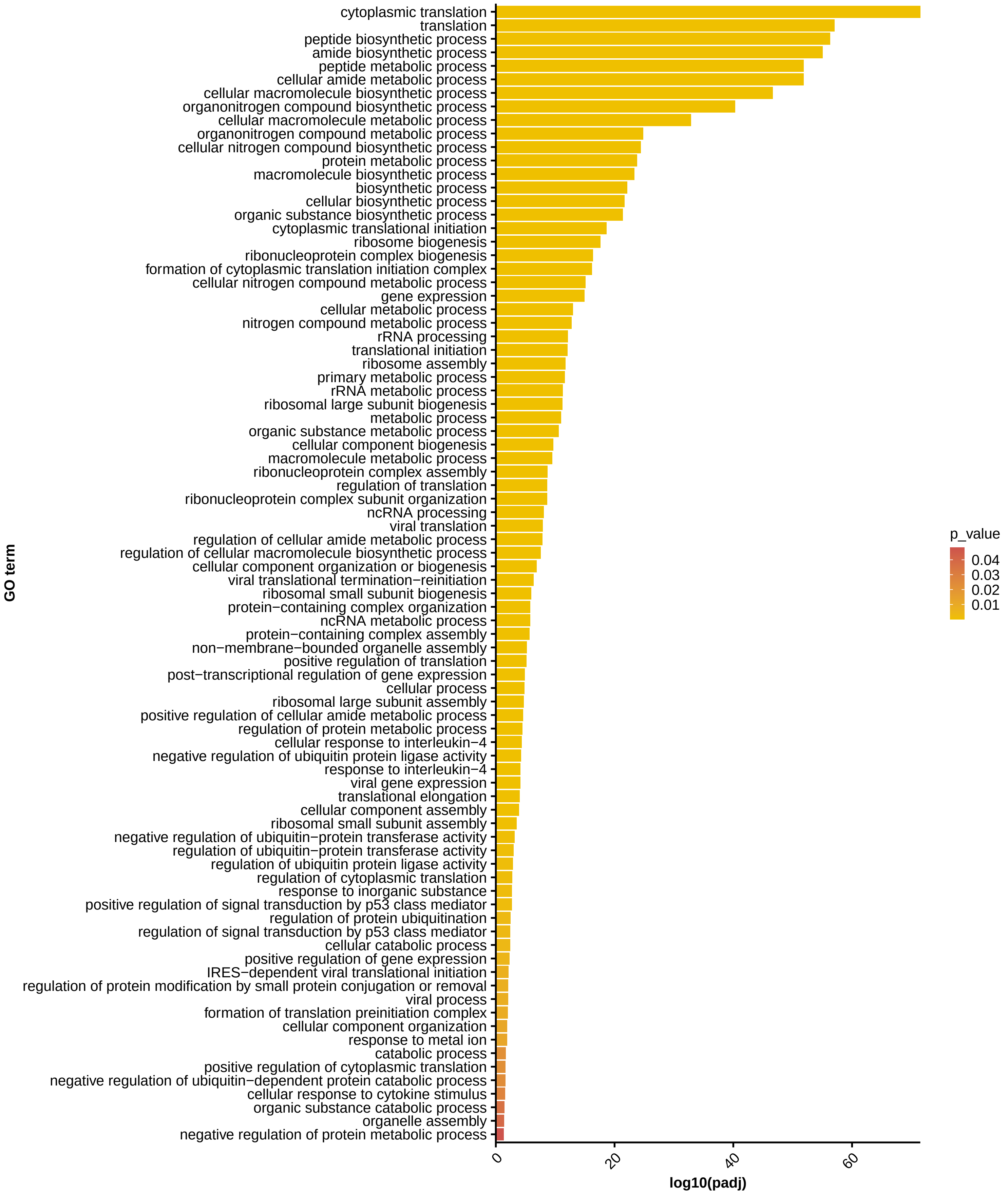

**Supplementary Figure 3.** Gene Ontology Biological Proces terms identified in functional annotation analysis of proteins coprecipitated with DIS3L-GFP.

**Supplementary Figure 4**

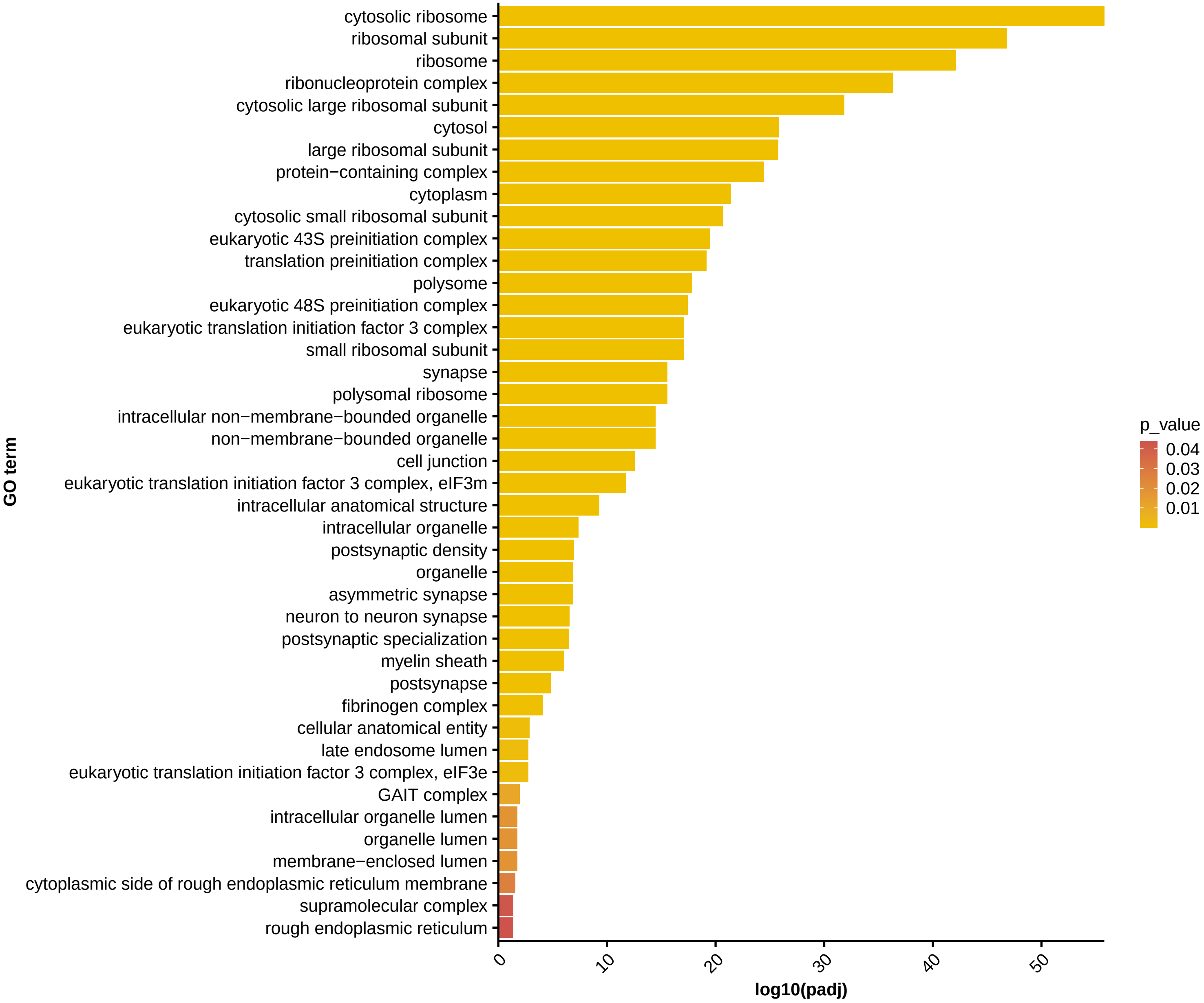

**Supplementary Figure 4.** Gene Ontology Cellular Component terms identified in functional annotation analysis of proteins coprecipitated with DIS3L-GFP.
